# Localized phosphorylation of RNA Polymerase II by G1 cyclin-Cdk promotes cell cycle entry

**DOI:** 10.1101/2021.03.25.436872

**Authors:** Mardo Kõivomägi, Matthew P. Swaffer, Jonathan J. Turner, Georgi Marinov, Jan M. Skotheim

**Affiliations:** Department of Biology, Stanford University, Stanford CA 94305, USA; Department of Genetics, Stanford University, Stanford CA, 94305, USA

## Abstract

The cell cycle is thought to be initiated by cyclin-dependent kinases (Cdk) inactivating transcriptional inhibitors of cell cycle gene-expression*(1, 2)*. In budding yeast, the G1 cyclin Cln3-Cdk1 complex is thought to directly phosphorylate Whi5, thereby releasing the transcription factor SBF and committing cells to division*(3-7)*. Here, we report that Cln3-Cdk1 does not phosphorylate Whi5, but instead phosphorylates the RNA Polymerase II subunit Rpb1’s <u>C</u>-<u>t</u>erminal <u>d</u>omain (CTD) on S_5_ of its heptapeptide repeats. Cln3-Cdk1 binds SBF-regulated promoters*(8)* and Cln3’s function can be performed by the canonical S_5_ kinase*(9)* Ccl1-Kin28 when synthetically recruited to SBF. Thus, Cln3-Cdk1 triggers cell division by phosphorylating Rpb1 at SBF-regulated promoters to activate transcription. Our findings blur the distinction between cell cycle and transcriptional Cdks to highlight the ancient relationship between these processes.

---

The eukaryotic cell cycle is driven by a series of cyclin-Cdk complexes that promote cell cycle progression by phosphorylating key substrates*(2)*. The first step of the eukaryotic cell cycle, from G1 to S phase, has long been thought to require the phosphorylation and inactivation of a transcriptional inhibitor. In human cells, cyclin D-Cdk4,6 phosphorylates the retinoblastoma protein, Rb, and in budding yeast Cln3-Cdk1 is thought to phosphorylate Whi5*(1, 3, 4)*. This results in the activation of the E2F and SBF transcription factors, in animal and yeast cells respectively, which then commits cells to division via positive feedback loops (Fig. 1A)*(5, 10-15)*. However, in early to mid G1, cyclin D-Cdk4,6 constitutively hypo-phosphorylates Rb*(16)*, which is likely insufficient to completely inactivate Rb*(17)*. As for Whi5, phosphorylation by Cln3-Cdk1 has not previously been observed *in vivo*. Thus, key mechanistic aspects of the eukaryotic G1/S transition model remain either unknown or untested.

**Fig. 1.**
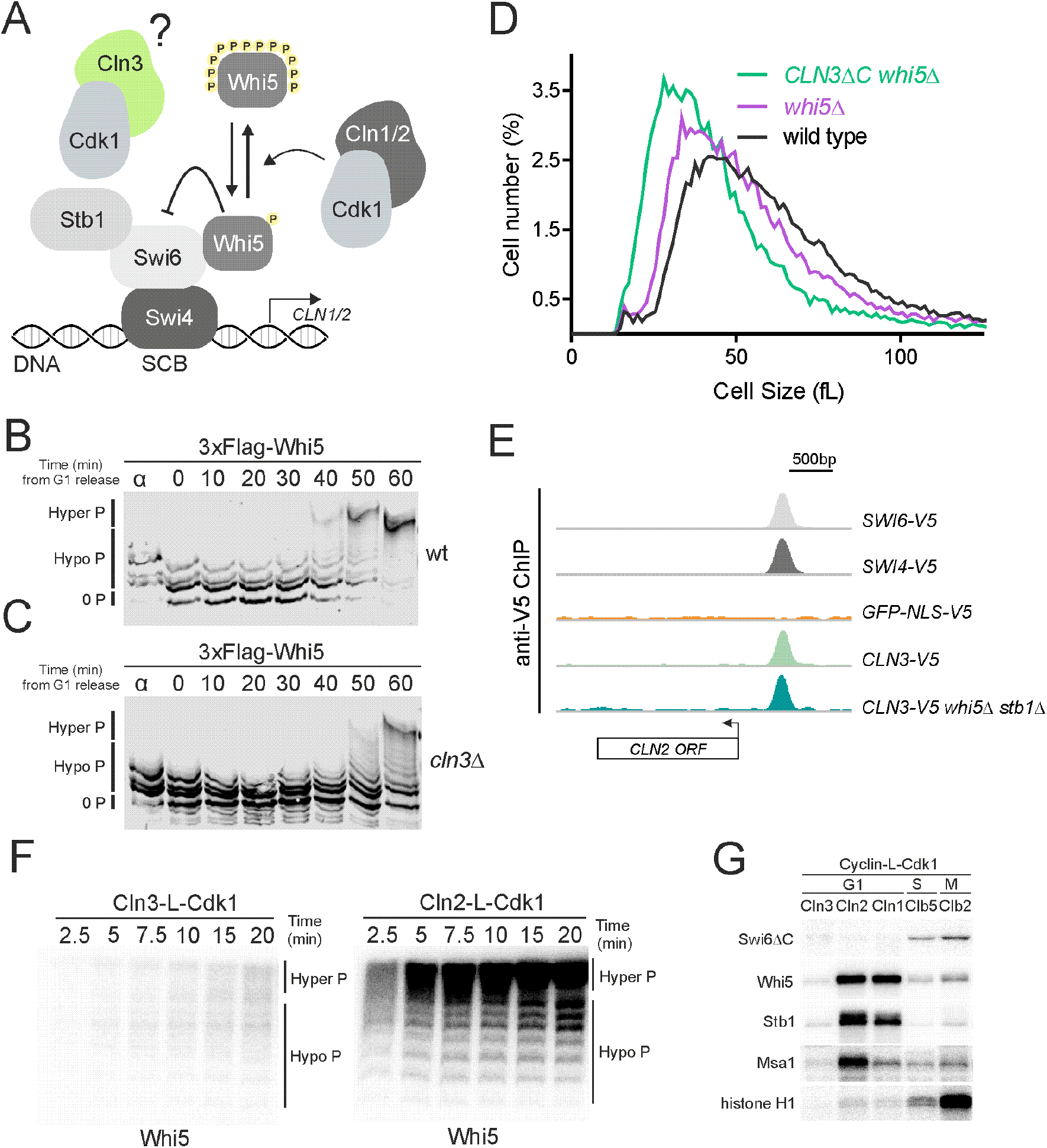
Cln3-Cdk1 binds SBF, but does not phosphorylate the transcriptional inhibitor Whi5. (**A**) Schematic of the budding yeast G1/S regulatory network. (**B**) Phos-tag immunoblot time course measuring distinct hypo- and hyper-phosphorylated isoforms of 3xFlag-Whi5 after release from a G1 pheromone arrest. (**C**) Phos-tag immunoblot time course as in (B) for *cln3*Δ cells. (**D**) Cell size distributions measured by Coulter counter for the indicated genotypes. Cells were grown on synthetic complete media with 2% glucose. (**E**) anti-V5 ChIP-seq signal of the indicated genotypes at the *CLN2* locus. *CLN2* is an SBF target, whose expression drives the G1/S transition. See fig. S1E to F and methods for more details. (**F**) Autoradiographs of *in vitro* Whi5 phosphorylation time courses by Cln2-L-Cdk1 and Cln3-L-Cdk1 fusion proteins, where L denotes a glycine serine linker (see methods for purification and verification of Cln3-L-Cdk1 activity; fig. S3C). Whi5 phospho-isoforms were resolved using Phos-tag SDS-PAGE. All reactions contain equal amounts of the indicated cyclin-L-Cdk1 complexes. (**G**) Autoradiographs of *in vitro* phosphorylation of SBF-interacting proteins and histone H1 by the indicated cyclin-L-Cdk1 complexes. All reactions contain equal amounts of the respective cyclin-L-Cdk1 complexes.

To examine the prevailing model that Cln3-Cdk1 promotes the G1/S transition by progressively phosphorylating and inhibiting Whi5*(3, 4)*, we sought to measure Whi5 phosphorylation *in vivo* as cells progressed through G1. We used Phos-tag-supplemented SDS-PAGE*(18)* to separate distinct phospho-isoforms of Whi5 isolated from cells synchronously released from G1 arrest. This allowed us to resolve not only multi-phosphorylated species of Whi5 but also different mono- or di-phosphorylated species, which had never previously been observed (Fig. 1B). Hypophosphorylation of Whi5 is slightly reduced upon release from pheromone arrest, but stays at a constant level until the G1/S transition. At G1/S, Whi5 is rapidly hyper-phosphorylated by Cln1/2-Cdk1*(3, 6)*.

Next, we sought to test if Whi5 hypophosphorylation in early G1 is due to Cln3-Cdk1. Consistent with Cln3’s well-established role in driving G1/S, Whi5 hyperphosphorylation was delayed in *cln3*Δ cells synchronously released into the cell cycle as previously reported*(19)* (Fig. 1C). But, critically, the hypophosphorylation pattern was the same as in WT cells in early-to mid-G1, a time when Cln3 is constitutively expressed and thought to function*(7, 20)*. Furthermore, the hypophosphorylation pattern was unaffected by conditionally expressing *CLN3* in G1-arrested *cln3*Δ*bck2*Δ cells, or by inhibiting an ATP analog-sensitive Cdk1 (Cdk1^as^) in mid-G1 (fig. S1A to C). In contrast, inhibiting Cdk1^as^ in S/G2, after Whi5 was already hyperphosphorylated, caused rapid Whi5 dephosphorylation back to the same G1 hypophosphorylated isoforms (fig. S1D). Taken together, these experiments argue strongly against the prevailing model that Cln3-Cdk1 phosphorylates Whi5 to drive G1/S. Rather, they raise the possibility that Cln3 and Whi5 act as separate inputs regulating SBF activity. Consistent with this separate input model, *cln3*Δ*whi5*Δ cells were larger than *whi5*Δ cells*(3, 4)*, and addition of a hyperactive *CLN3* allele *(CLN3*Δ*C)(21, 22)* reduced cell size more than *whi5*Δ (Fig. 1D).

If Cln3-Cdk1 functions through SBF but does not target Whi5, it might be present at SBF-regulated promoters even in the absence of Whi5. To test this, we performed ChIP-seq analysis of Cln3 and the SBF components Swi4 and Swi6, all tagged at their endogenous loci with the V5 epitope. Cln3-V5 was found at 85 gene promoters, 84 of which were also bound by SBF (Swi4-V5 and Swi6-V5; Table S3). These sites include key SBF binding-sites in the *CLN1* and *CLN2* promoters (Fig. 1E, fig. S1E to F), consistent with previous ChIP experiments showing conditionally-expressed Cln3 binding at the *CLN2* promoter*(8)*. Furthermore, Cln3-V5 localization to SBF-binding sites, including the *CLN2* promoter, did not depend on Stb1 or Whi5, showing that the presence of Cln3 at SBF sites did not depend on its previously assumed target protein (Fig. 1E, fig. S1E to F).

That Cln3 binds SBF-regulated promoters suggests that Cln3-Cdk1 phosphorylates a different target involved in SBF-dependent transcription. To explore this possibility, we examined Cln3-Cdk1 kinase activity towards SBF-interacting proteins *in vitro*. To purify Cln3-Cdk1, we fused *CLN3* to *CDK1* using a glycine-serine linker*(23, 24) (CLN3-L-CDK1)* because Cln3 is not as tightly bound to Cdk1 as the other yeast cyclins. When the endogenous *CLN3* allele was replaced by this fusion allele, cells exhibited no cell cycle defects and were the same size as wild type, suggesting that this fusion complex functions similarly to the wild type allele (fig. S2A). We note that, to the best of our knowledge, this is the first time active Cln3-Cdk1 complexes have been purified to homogeneity. Kinase activity detected in a previously reported purification of Cln3-Cdk1 expressed in insect cells*(4)*, which we repeated, was not due to Cln3 because this activity was still present in a negative control lacking Cln3 (see methods; fig. S2B). Consistent with our *in vivo* phosphorylation data, Cln3-L-Cdk1 poorly phosphorylated Whi5 *in vitro*, while Cln2-L-Cdk1 readily hyperphosphorylated Whi5 (Fig. 1F). Moreover, Cln3-L-Cdk1 poorly phosphorylated the SBF-associated proteins*(8, 25-27)* Swi6, Stb1, and Msa1 *in vitro* (Fig. 1G). This is consistent with a previous study concluding that although the SBF subunit Swi6 was required for Cln3 function, it was not a direct target of Cln3-Cdk1*(28)*.

The absence of Cln3-dependent phosphorylation on proteins at SBF-regulated promoters, combined with Cln3-Cdk1’s lack of *in vitro* activity against the model Cdk substrate H1 (Fig. 1G), suggested that Cln3 may promote the G1/S transition independently of Cdk1 kinase activity. To test this possibility, we examined the effect of replacing *CLN3* with a *CLN3* allele fused to a previously described kinase dead *CDK1* allele*(29)(CDK1*^*KD*^; Fig. 2A, fig. S2C). Although *CLN3-L-CDK1*^*KD*^ rescued the effects of *cln3*Δ, immunoprecipitation revealed endogenous Cdk1 bound to the fusion protein (fig. S2D). To prevent this, we introduced a <u>c</u>yclin <u>b</u>ox <u>m</u>utation to *CLN3* that prevents its binding to Cdk1*(30) (CLN3*^*CBM*^; Fig. 2A). Replacing *CLN3* with *CLN3*^*CBM*^ resulted in a similar large size phenotype as *cln3*Δ, while replacing *CLN3* with *CLN3*^*CBM*^*-L-CDK1* resulted in wild type sized cells, indicating that the fusion allows Cln3^CBM^ to activate Cdk1 (Fig. 2B). *CLN3*^*CBM*^*-L-CDK1*^*KD*^ cells lacking Cln3 activity were similar in size to *cln3*Δ cells, suggesting that Cln3-Cdk1 kinase activity is indeed required to drive the cell cycle through G1/S (Fig. 2B).

**Fig. 2.**
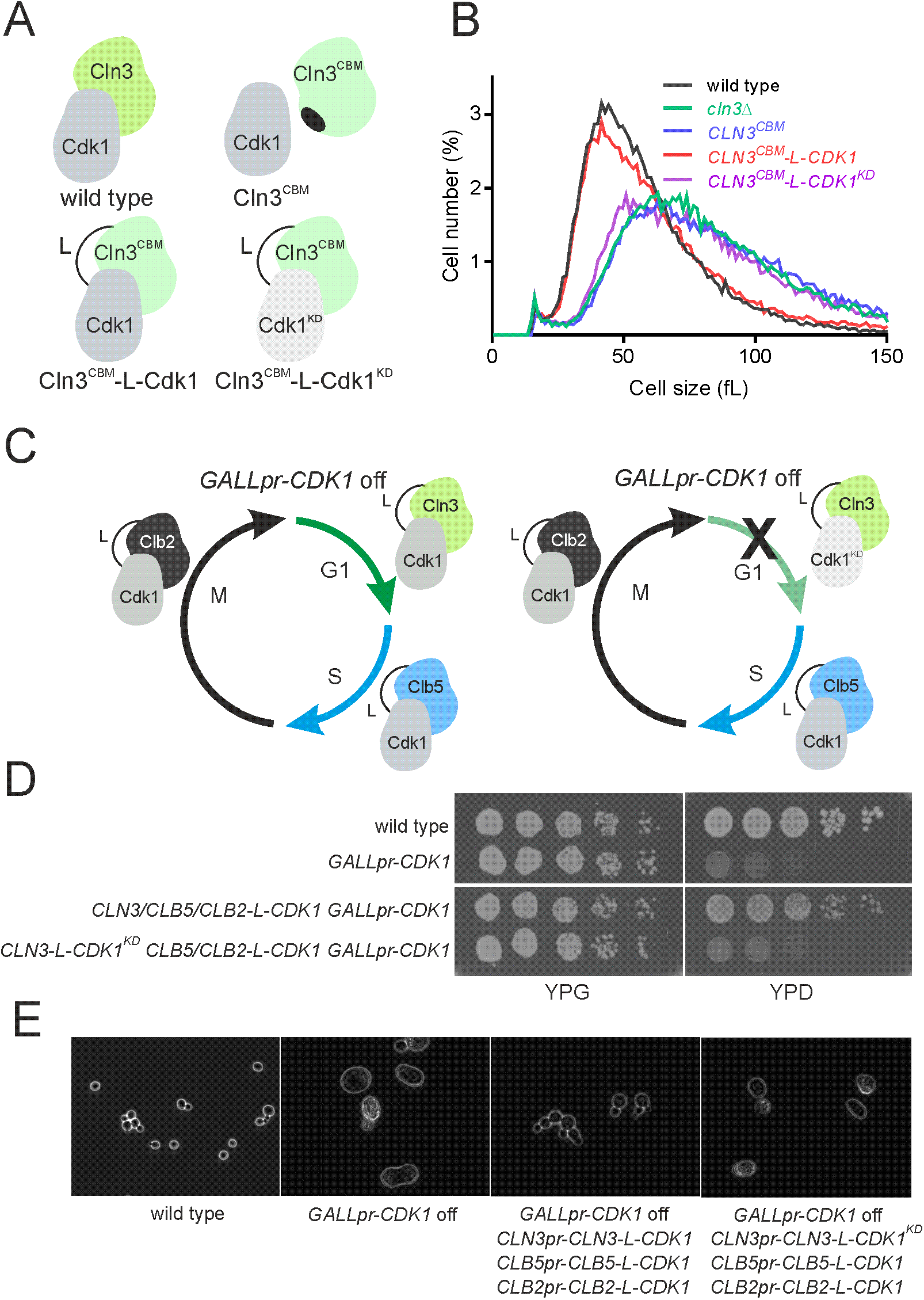
Cln3-Cdk1 kinase activity promotes the G1/S transition. (**A**) Schematic of the different Cln3-Cdk1 complexes used. Cln3^CBM^ denotes a <u>c</u>yclin <u>b</u>ox <u>m</u>utant that does not bind Cdk1 unless fused via the linker, L. Cdk1^KD^ denotes a <u>k</u>inase <u>d</u>ead mutant of Cdk1. (**B**) Cell size distributions measured by Coulter counter for the indicated genotypes. Cells were grown on synthetic complete media +2% glucose. (**C**) Schematic of the experimental design shown in (D). The endogenous *CDK1* promoter was replaced with the galactose-inducible *GALL* promoter *(GALLpr)*. *CLN3, CLB5*, and *CLB2* were then fused at their endogenous locus to *CDK1*. (**D**) Spot viability assays of WT and *GALLpr-CDK1* strains the on YPG *(GALLpr* ON) or YPD *(GALLpr* OFF). The triple *cyclin-CDK1* fusion rescues *CDK1* repression only if *CLN3* is fused to an active *CDK1*. (**E**) Phase contrast images of cells of the indicated genotypes after *GALLpr-CDK1* repression.

To confirm that Cln3-Cdk1 requires kinase activity to promote cell-cycle progression, we examined its function in the context of a simplified cell-cycle control network. We first replaced the *CDK1* promoter with the glucose-repressible *GAL1* promoter so that all endogenous Cdk1 activity could be conditionally removed. We then proceeded to add back cyclin-Cdk1 fusion proteins expressed from their endogenous cyclin promoters. Cells were viable on glucose when *CLB2-L-CDK1* was added to drive M-phase, *CLB5-L-CDK1* was added to drive S-phase, and *CLN3-L-CDK1* was added to drive G1/S (Fig. 2C to E). However, addition of *CLN3-L-CDK1*^*KD*^ instead of *CLN3-L-CDK1* was insufficient for proliferation, which supports a model in which Cln3-Cdk1 kinase activity promotes the G1/S transition (Fig. 2C to E). The requirement for Cln3-Cdk1 activity was not alleviated by deletion of *WHI5*, which is consistent with Whi5 not being a Cln3-Cdk1 substrate (fig. S2F to G).

Having established that Cln3-Cdk1 requires kinase activity to promote the G1/S transition, we sought to identify its substrates. We performed a candidate-based *in vitro* screen*(31)*, in which we measured the activity of purified Cln3-L-Cdk1 and other yeast cyclin-Cdk1 complexes towards >20 Cdk1 target proteins (Fig. 3A to C; fig. S3A to D). By far the most specific target for Cln3-L-Cdk1 was the RNA polymerase II subunit Rbp1, which contains a C-terminal unstructured region (CTD) with multiple heptapeptide repeats (26 in yeast, 52 in humans) of the sequence Y_1_S_2_P_3_T_4_S_5_P_6_S_7_ *(32)* (Fig. 3C). Truncations of Rpb1 to first isolate the unstructured C-terminal region and then to remove the regions on either side of the CTD heptad repeats did not reduce phosphorylation, implying that Cln3-Cdk1 directly targets one or multiple residues inside the heptapeptide repeats independently of the adjacent unstructured regions (Fig. 3D to E).

**Fig. 3.**
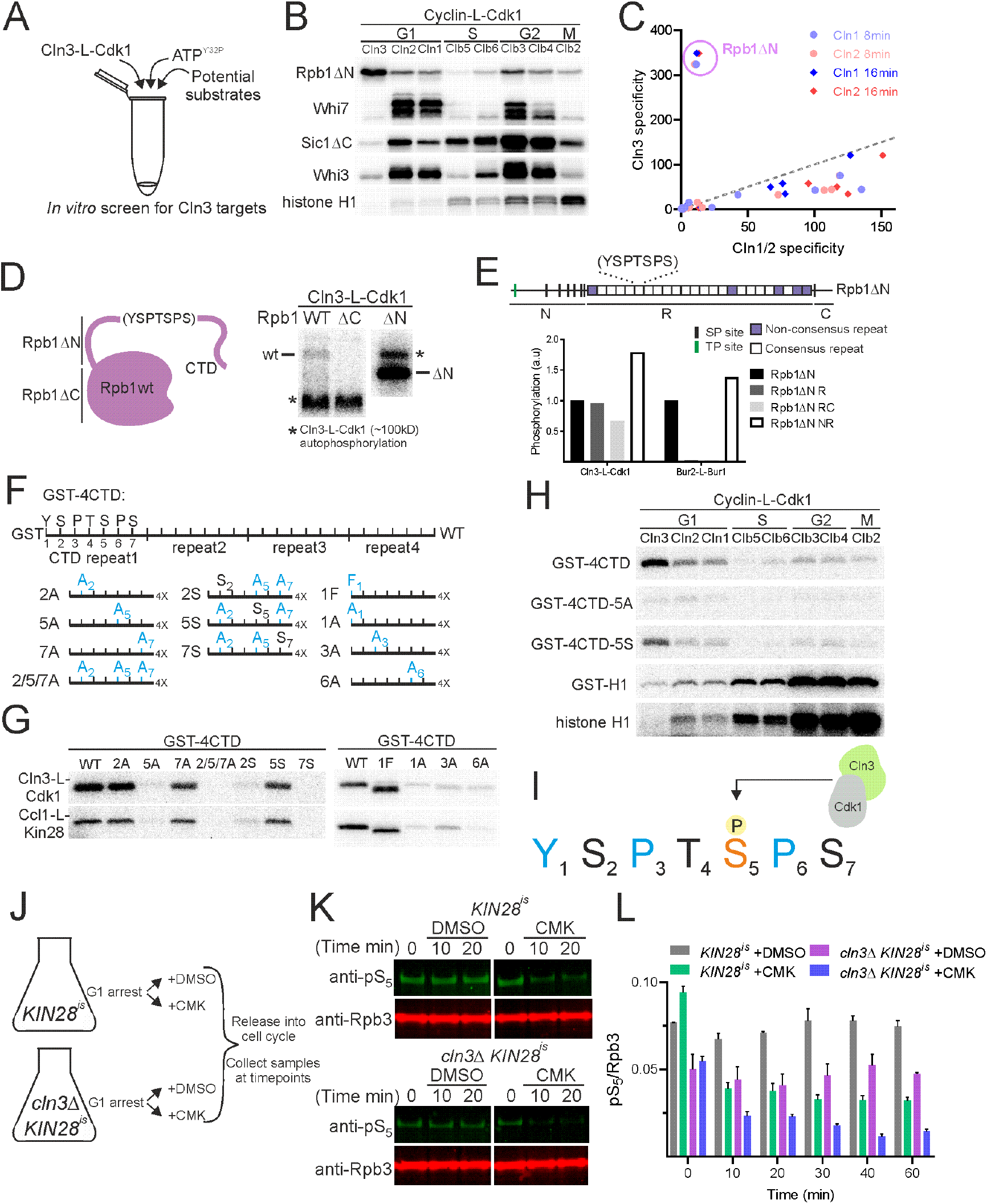
Cln3-Cdk1 phosphorylates the serine 5 residue in the Pol II subunit Rpb1’s C-terminal domain repeats. (**A**) Schematic of *in vitro* candidate based screen for Cln3-Cdk1 targets. (**B**) Autoradiographs of *in vitro* phosphorylation of a subset of candidate substrates by the indicated cyclin-L-Cdk1 complexes. All reactions contain equal amounts of the indicated cyclin-L-Cdk1 complexes. (**C**) Quantification of *in vitro* phosphorylation specificity by Cln3-L-Cdk1 compared to Cln1-L-Cdk1 or Cln2-L-Cdk1. Specificity is defined as the ratio of activity towards the indicated substrate relative to the activity towards the Cdk1 model substrate histone H1. Each point corresponds to a single substrate (n=20) and quantifications from two time points are plotted. Dotted line denotes x=y where Cln3-L-Cdk1 substrate specificity is equally to Cln1-L-Cdk1 or Cln2-L-Cdk1 specificity. See methods and Table S4 for measurements. (**D**) Autoradiographs of *in vitro* phosphorylation by Cln3-L-Cdk1 of Rpb1 and Rpb1 truncations (WT denotes full length; ΔC denotes the 1453 N-terminal residues; ΔN denotes the 280 C-terminal residues). We note the low yield of full length Rpb1 (∼192kD) results in a lower phosphorylation signal than the ΔN. Full length and ΔC Rpb1 were resolved on 6% SDS PAGE gels, while ΔN was resolved on a separate 10% SDS PAGE gel. (**E**) Autoradiograph quantification from *in vitro* kinase assays phosphorylating Rpb1ΔN and Rpb1ΔN truncations by Cln3-L-Cdk1 and the transcriptional kinase Bur2-L-Bur1. See fig. S3G for autoradiograph images including data for other transcriptional kinases. (**F**) Schematic of GST epitope model substrates used in (G), containing 4 copies of the CTD repeat. (**G**) Autoradiographs of *in vitro* phosphorylation of CTD repeats with the indicated amino acid substitutions by Cln3-L-Cdk1 and Ccl1-L-Kin28. (**I**) Schematic showing S_5_ specific phosphorylation by Cln3-Cdk1. (**J**) Schematic of experimental design for (K-L). *Kin28*^*is*^ or *Kin28*^*is*^ *cln3*Δ cells were released from G1 pheromone arrest into DMSO or CMK. *Kin28*^*is*^ contains an active site mutant rendering it sensitive to covalent inhibition by the small molecule CMK. (**K**) Representative immunoblots for total cellular phosphorylated Rpb1-CTD S_5_ (H14 antibody) and total cellular RNA polymerase II (Rpb3) after release from G1 pheromone arrest into DMSO or CMK. See fig. S4C for full immunoblots. (**L**) Quantification of total cellular phosphorylated Rpb1-CTD S_5_ (H14 antibody) normalized to total cellular RNA polymerase II (Rpb3). Mean ± S.E.M. is plotted, calculated from two independent biological replicates.

Phosphorylation of the different residues within these heptad repeats by the canonical transcriptional kinases regulates transcriptional initiation, elongation, and termination*(32)*. To compare Cln3-Cdk1 with the four known transcriptional kinases, we applied our *in vitro* approach to purify Ccl1-L-Kin28, Bur2-L-Bur1, Ctk2-L-Ctk1 and Ssn8-L-Ssn3, which correspond to human Cdk7, Cdk9, Cdk12, and Cdk8 complexes respectively*(33)*. This revealed Cln3-Cdk1, Kin28, Ssn3 and Ctk1 all phosphorylate residues inside the CTD repeats independently of the adjacent unstructured regions (fig. S3E to G). In contrast, Bur1 did not phosphorylate the CTD repeats without the C-terminal non-repeat element (Fig. 3E), consistent with published work*(34)*.

To determine the function of Cln3-Cdk1 phosphorylation of the CTD, we sought to identify the specific target residues. To do this, we generated a series of model substrates comprising a GST sequence followed by four wild type or mutant CTD consensus repeats (GST-4CTD) (Fig. 3F). Of the possible Cdk phosphorylation site mutants, only mutation of the serine 5 residue prevented phosphorylation (GST-4CTD_5A_). Conversely, the addition of S_5_ to a repeat region lacking all serines restored phosphorylation (GST-4CTD_S5_ in Fig. 3G). Similar results were found when this set of substrates was phosphorylated by Ccl1-L-Kin28, but not when phosphorylated by other cyclin-Cdk1 complexes (Fig. 3G to H). That Cln3-Cdk1 could function as an S_5_ CTD kinase is consistent with the reported genetic interactions between the canonical S_5_ CTD kinase Kin28 and Cdk1*(35)*. To determine if other residues inside the CTD heptad might be responsible for Cln3-Cdk1 specificity, we performed *in vitro* kinase assays with additional GST-4CTD substrates in which Y_1_, P_3_, T_4_, or P_6_ were substituted with alanines. Phosphorylation was decreased by Y_1_, P_3_, and P_6_ mutations, but not by T_4_ mutation (Fig. 3G; fig. S3E). The effect of the P_6_ alanine substitution was expected because Cdk1 is a proline directed kinase, while the P_3_ requirement 2 residues N-terminal to the phosphorylation site is similar to that found to enhance phosphorylation by Cln2-Cdk1 complexes*(31)*. However, the Y_1_ requirement was surprising and suggests a highly specific substrate preference for the active site of Cln3-Cdk1. Taken together, our results show that Cln3-Cdk1 likely phosphorylates S_5_ and that this phosphorylation depends on the local amino acid sequence (Fig. 3I).

That Cln3 localizes to specific SBF-regulated promoters, and that Cln3-L-Cdk1 functions as an S_5_ CTD kinase *in vitro*, suggests a model in which Cln3-Cdk1 promotes transcription of SBF-regulated genes by phosphorylating the CTD of Rpb1 at their promoters. In this model, Cln3-Cdk1 should be responsible for only a subset of the global S_5_ CTD phosphorylation. To test this model, we first replaced endogenous *KIN28* with the *KIN28*^*is*^ allele, which expresses a version of Kin28 that can be irreversibly inhibited by a covalently-binding small molecule*(36)*. Addition of the Kin28 inhibitor reduced S_5_ CTD phosphorylation by 60±3%, and deletion of *CLN3* further reduced phosphorylation by another 15±5% in cells in the G1 phase of the cell cycle, showing that Cln3-Cdk1 phosphorylates S_5_ in the CTD repeats *in vivo* (Fig. 3J to L, fig. S4; see methods).

Taken together, our work supports a model in which Cln3-Cdk1 promotes the G1/S transition by phosphorylating S_5_ in the RNA polymerase II CTD at SBF-regulated promoters (Fig. 4A). This model predicts that we should be able to bypass the requirement for Cln3 by providing an alternative source of S_5_ phosphorylation to SBF-regulated promoters. To test this, we used a rapamycin-dependent inducible binding system to conditionally recruit a fusion protein of the canonical CTD S_5_ kinase Ccl1-L-Kin28 to SBF via its Swi6 or its Swi4 subunit (Fig. 4B, fig. S5A and B). We note that all these strains contain the *tor1-1 fpr1*Δ mutations so that growth is not affected by rapamycin*(37)*. Strikingly, recruitment of Ccl1-L-Kin28 to SBF fully rescues the size and cell cycle phenotypes of *cln3*Δ cells and this rescue is dependent on Kin28 kinase activity (Fig. 4C to E, fig. S5C to F). This rescue was not due to Kin28 phosphorylation of Whi5 (fig. S5G to J).

**Fig. 4.**
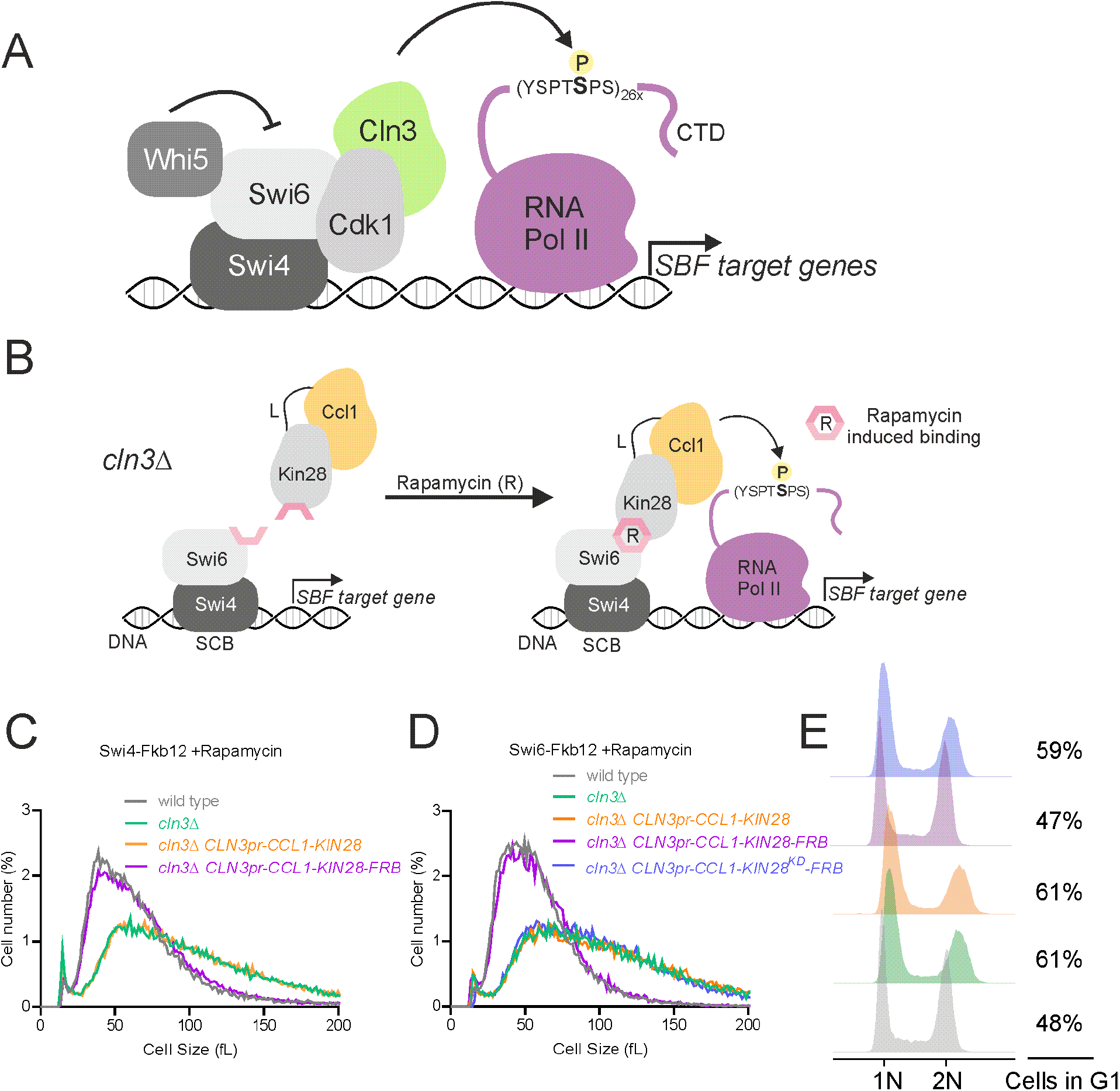
Induced binding of the S_5_ Rpb1 CTD kinase Ccl1-Kin28 to SBF rescues the cell cycle defects of *cln3*Δ cells. (**A**) Model in which Cln3-Cdk1 activates transcription at SBF-dependent promoters by phosphorylating Rpb1’s CTD on S_5_. (**B**) Schematic of experiment in (c-e): conditional recruitment of Ccl1-L-Kin28 to SBF using the rapamycin inducible binding system. (**C-D**) Cell size distributions measured by Coulter counter for the indicated genotype. Ccl1-L-Kin28 fusion proteins were expressed from a genomically integrated copy of the *CLN3* promoter. Ccl1-L-Kin28-FRB was recruited to SBF via Swi4 (C) or Swi6 (D) upon rapamycin treatment. Ccl1-L-Kin28 lacking FRB is not recruited. All strains were grown on synthetic complete media with 2% glucose and with 1µg/ml rapamycin or DMSO. *KIN28*^*KD*^ denotes a kinase-dead *KIN28*, whose recruitment to SBF does not rescue defects associated with *cln3*Δ. (**E**) Flow cytometry analysis of DNA content of the cells in (D).

Thus, Cln3-Cdk1 promotes the first step in the budding yeast cell cycle by directly phosphorylating RNA Polymerase II at SBF-regulated genes. Surprisingly, Cln3-Cdk1 did not phosphorylate its expected target, the SBF inhibitor Whi5, or any other proteins on the SBF transcription factor complex. Moreover, our screen suggests that Cln3-Cdk1 has few, if any, targets other than the RNA Polymerase II subunit Rpb1’s CTD. Cln3-Cdk1’s extreme specificity—for a single target driving the first step of the cell cycle—may help order cell cycle events*(31)*. If, as we suspect, Cln3 has no targets driving replication, spindle pole body duplication, cell polarization, or any other cell cycle event, the level of Cln3 can be used to exclusively modulate the G1/S transition without risking premature triggering of downstream cell cycle events.

That Cln3 and Whi5 serve as separate inputs to SBF activity can be rationalized by their potentially separate functions. In G1, Whi5 concentration directly reflects cell size because Whi5 is a stable protein and a constant number of Whi5 molecules is synthesized in S/G2/M phases independent of cell size and growth conditions*(20, 38)*. As cells grow in G1, Whi5 is then diluted so that its concentration is determined by cell size. In contrast, Cln3 concentration is constant in G1 as cells grow in a given condition, but this constant G1 concentration is higher when cells are growing rapidly, as in glucose, and lower, when cells are growing more slowly, as in ethanol*(7, 39-43)*. Thus, Cln3 and Whi5 may independently reflect cell growth and size, respectively.

While the cell cycle Cdks and the transcriptional Cdks are all part of the same kinase family, their functions are largely thought to have diverged along their separate branches*(44, 45)*. Our work here breaks down the previous dichotomy of cell cycle and transcriptional Cdks and shows how cell cycle Cdks can directly activate transcription at specific target genes to drive a cell cycle transition. The functional overlap of cell cycle and transcriptional Cdks has already been pointed to by the dual function of the Kin28 orthologs, Msc6 and Cdk7 in fission yeast and vertebrate cells, respectively. They both activate the cell cycle Cdks by phosphorylating their T-loops while also activating transcription as a global S_5_ CTD kinase*(46-48)*. In addition, Cdk1 and Cdk2 were identified as the first RNA Pol II CTD kinases *in vitro*, but if they function in such a manner *in vivo* remains unknown*(49)*. That the two branches of Cdks that regulate cell division and transcription have overlapping functions suggests the possibility that their primordial ancestor regulated both processes. These functions would then have been partially lost along the two divergent branches. Thus, our discovery that yeast Cln3-Cdk1 drives cell cycle progression by directly activating transcription may reflect an ancient link between basic biosynthetic processes like transcription and the control of cell division.

## TABLES AND TABLE CAPTIONS

**Table S1:**
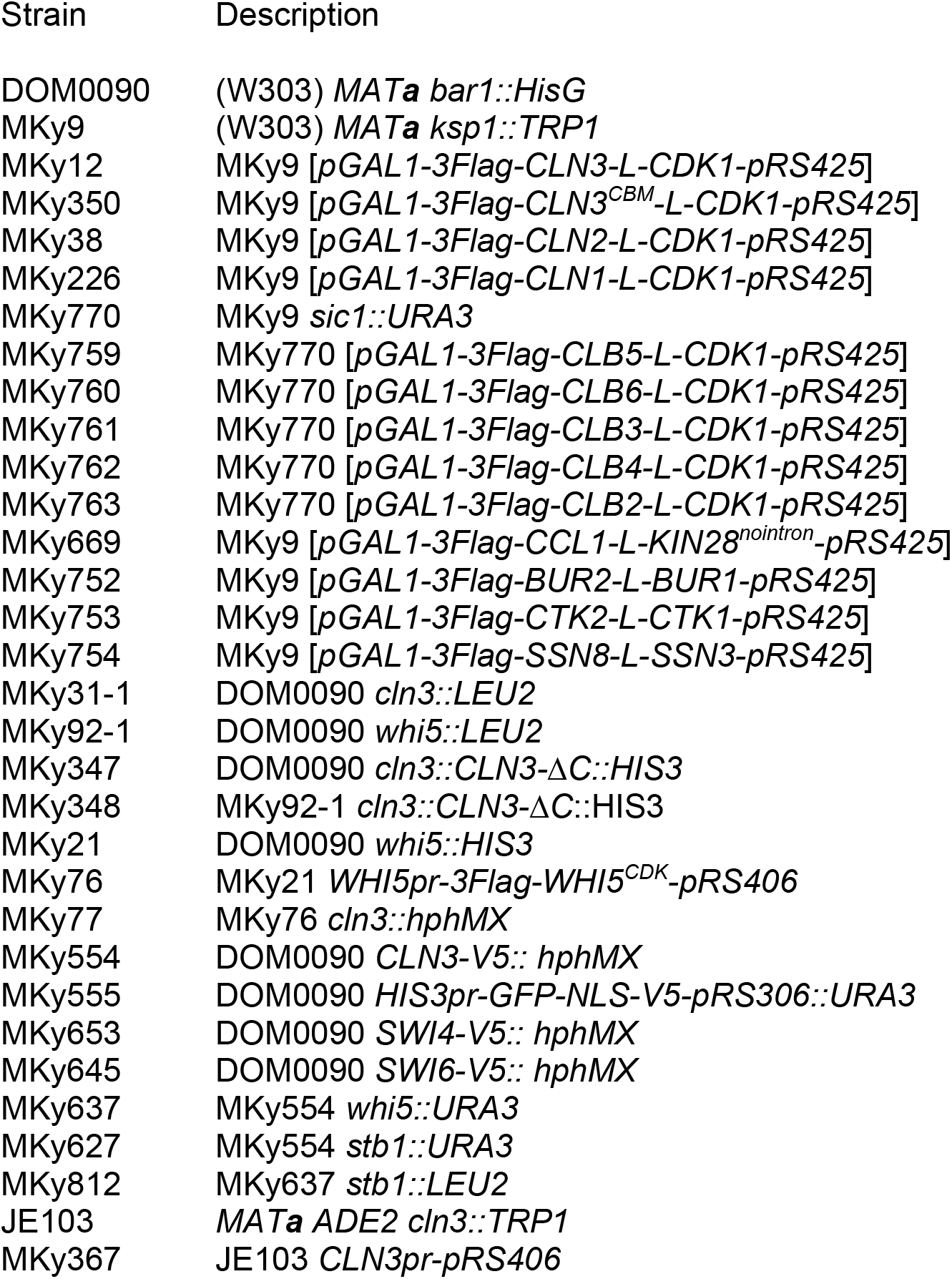
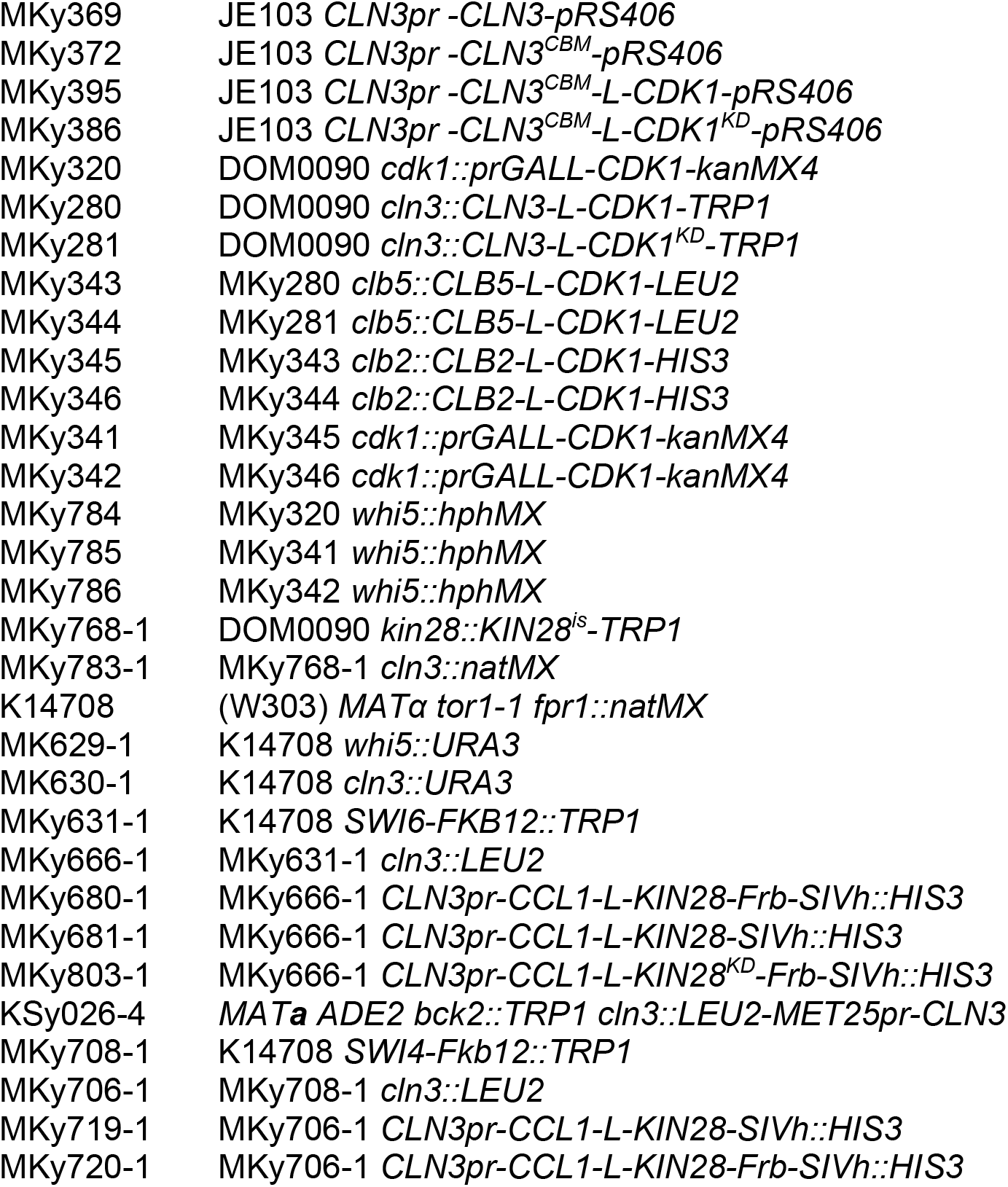
*Saccharomyces cerevisiae* strains used in this study. Parentheses denote plasmids, unless otherwise noted all other genotypes represent genomic integration. Strains and plasmids were made using standard methods. DOM0090 was obtained from David Morgan, JE103 was obtained from Jennifer Ewald*(50)*, and K14708 was obtained from Euroscarf*(37)*. *CLN3*^*CBM*^ denotes the mutations to the cyclin box of *CLN3(30)*. *KIN28*^*nointron*^ denotes an allele of *KIN28* where the intron was removed. *CLN3-*Δ*C* denotes an allele of *CLN3* lacking the C-terminal unstructured region, *i*.*e*., the allele is truncated at bp1197. *WHI5*^*CDK*^ denotes an allele of *WHI5* where two non-Cdk1 sites that we found to be phosphorylated in G1 were removed (Kõivomägi et al *in preparation)*. *CDK1*^*KD*^ denotes a kinase dead allele of *CDK1* where K40 was mutated to L*(29)*. *KIN28*^*is*^ denotes a *KIN28* allele that can be covalently inhibited*(36)*. *SIC1dC* denotes a *SIC1* allele truncated at bp645*(51)*. *RPB1dN* (bp4360-5202) and *RPB1dC* (bp1-4359) denote truncations of either the N- or C-terminal unstructured regions. Subscript numbers denote the basepairs present of the gene indicated that were present.

**Table S2:** Plasmids used in this study.

|  |  |  |
| --- | --- | --- |
| 386 | Plasmid | Description |
| 387 | pMK178 | *pGAL1-3Flag-CLN3-L-CDK1-pRS425* |
| 388 | pMK185 | *pGAL1-3Flag-CLN3<sup>CBM</sup>-L-CDK1-pRS425* |
| 389 | pMK184 | *pGAL1-3Flag-CLN2-L-CDK1-pRS425* |
| 390 | pMK186 | *pGAL1-3Flag-CLN1-L-CDK1-pRS425* |
| 391 | pMK361 | *pGAL1-3Flag-CLB5-L-CDK1-pRS425* |
| 392 | pMK362 | *pGAL1-3Flag-CLB6-L-CDK1-pRS425* |
| 393 | pMK362 | *pGAL1-3Flag-CLB3-L-CDK1-pRS425* |
| 394 | pMK364 | *pGAL1-3Flag-CLB4-L-CDK1-pRS425* |
| 395 | pMK366 | *pGAL1-3Flag-CLB2-L-CDK1-pRS425* |
| 396 | pMK308 | *pGAL1-3Flag-CCL1-L-KIN28<sup>nointron</sup>-pRS425* |
| 397 | pMK367 | *pGAL1-3Flag-BUR2-L-BUR1-pRS425* |
| 398 | pMK368 | *pGAL1-3Flag-CTK2-L-CTK1-pRS425* |
| 399 | pMK369 | *pGAL1-3Flag-SSN8-L-SSN3-pRS425* |
| 400 | pMK187 | *WHI5pr-3Flag-WHI5<sup>CDK</sup>-pRS406* |
| 401 | pMK188 | *CLN3pr-pRS406* |
| 402 | pMK189 | *CLN3pr-CLN3-pRS406* |
| 403 | pMK190 | *CLN3pr-CLN3<sup>CBM</sup>-pRS406* |
| 404 | pMK191 | *CLN3pr-CLN3<sup>CBM</sup>-L-CDK1-pRS406* |
| 405 | pMK192 | *CLN3pr-CLN3<sup>CBM</sup>-L-CDK1<sup>KD</sup>-pRS406* |
| 406 | pMK159 | *CLN3<sub>1214-1740</sub>-L-CDK1-pRS403* |
| 407 | pMK160 | *CLN3<sub>1214-1740</sub>-L-CDK1<sup>KD</sup>-pRS403* |
| 408 | pMK161 | *CLB5<sub>841-1305</sub>-L-CDK1-pRS403* |
| 409 | pMK162 | *CLB2<sub>721-1473</sub>-L-CDK1-pRS403* |
| 410 | pMK0150 | *SIC1dC-pET28a* |
| 411 | pMK163 | *WHI5-pET28a* |
| 412 | pMK300 | *BDP1-pET28a* |
| 413 | pMK0078 | *STB1-pET28a* |
| 414 | pMK140 | *WHI3-pET28a* |
| 415 | pMK141 | *SIC1-pET28a* |
| 416 | pMK142 | *CLN3-pET28* |
| 417 | pMK143 | *CLN2-pET28a* |
| 418 | pMK144 | *RPD3-pET28a* |
| 419 | pMK145 | *CIP1-pET28a* |
| 420 | pMK146 | *CLN2dN-pET28a* |
| 421 | pMK147 | *CLN3dN-pET28a* |
| 422 | pMK370 | *RPB1dN-pET28a* |
| 423 | pMK371 | *RPB1dN-R-pET28a* |
| 424 | pMK372 | *RPB1dN-NR-pET28a* |
| 425 | pMK373 | *RPB1dN-RC-pET28a* |
| 426 | pMK374 | *SPT5dN-pET28a* |
| 427 | pMK148 | *WHI7-pET28a* |
| 428 | pMK149 | *DOA1-pET28a* |
| 429 | pMK150 | *GCR1-pET28a* |
| 430 | pMK151 | *BOI2-pET28a* |
| 431 | pMK152 | *CDC48dN-pET28a* |
| 432 | pMK153 | *YMR147-pET28a* |
| 433 | pMK0086 | *MSA1-pET28a* |
| 434 | pMK165 | *GST-WHI5-pGEX-4T-1* |
| 435 | pMK375 | *GST-RPB1dN-pGEX-4T-1* |
| 436 | pMK376 | *GST-MED15-pGEX-4T-1* |
| 437 | pMK377 | *GST-SWI6-pGEX-4T-1* |
| 438 | pMK378 | *GST-SWI4-pGEX-4T-1* |
| 439 | pMK379 | *GST-RPB1-pGEX-4T-1* |
| 440 | pMK380 | *GST-RPB1dC-pGEX-4T-1* |
| 441 | pMK381 | *GST-4CTD-pGEX-4T-1* |
| 442 | pMK382 | *GST-4CTD-1A-pGEX-4T-1* |
| 443 | pMK383 | *GST-4CTD-1F-pGEX-4T-1* |
| 444 | pMK384 | *GST-4CTD-2A-pGEX-4T-1* |
| 445 | pMK385 | *GST-4CTD-3A-pGEX-4T-1* |
| 446 | pMK386 | *GST-4CTD-5A-pGEX-4T-1* |
| 447 | pMK387 | *GST-4CTD-6A-pGEX-4T-1* |
| 448 | pMK388 | *GST-4CTD-7A-pGEX-4T-1* |
| 449 | pMK389 | *GST-4CTD-2/5/7A-pGEX-4T-1* |
| 450 | pMK390 | *GST-4CTD-2S-pGEX-4T-1* |
| 451 | pMK358 | *CLN3pr-CCL1-L-KIN28-Frb-SIVh* |
| 452 | pMK359 | *CLN3pr-CCL1-L-KIN28-SIVh* |
| 453 | pMK360 | *CLN3pr-CCL1-L-KIN28<sup>KD</sup>-Frb-SIVh* |
| 454 | pMK315 | *CCL1pr-CCL1-L-KIN28-Frb-SIVh* |
| 455 | pMK314 | *CCL1pr-CCL1-L-KIN28-SIVh* |
| 456 | pMS121 | *HIS3pr-GFP-NLS-V5-pRS306* |
| 457 | pBTC054 | *GST-H1-pGEX-4T-1* |

**Table S3: List of gene promoters overlapping with Cln3-V5 ChIP-seq peaks and associated peak coordinates**. See Methods for details.

**Table S4: List of substrates and quantification of their phosphorylation by G1 cyclin-Cdk1 complexes**.

## METHODS

### Yeast strains and plasmids

Standard procedures were used for growth and genetic manipulation of *Saccharomyces cerevisiae*. Cells were grown at 30°C in yeast extract/peptone medium with 2% glucose (YPD) or 2% galactose (YPGal), or in synthetic complete medium with 2% glucose (SCD) or 2% raffinose or with 2% glycerol and 1% ethanol. All *S. cerevisiae* strains in this study are derived from the W303 background. Full genotypes of all strains used in this study are listed in Table S1. Plasmids used in this study are listed in Table S2. For strains to conditionally inactivate Kin28, the endogenous *KIN28* gene was replaced with its *KIN28*^*is*^ counterpart by allele replacement into *cln3*Δ or wild type backgrounds*(36)*. After recombination, replacement of *KIN28* with *KIN28*^*is*^ was screened by growing colonies on rich media (YPD) or media containing 5µM CMK (YPD+CMK). Colonies that displayed a growth defect on YPD+CMK, but not on YPD, were selected and genotyped. For strains to conditionally recruit the Ccl1-L-Kin28 fusion protein to SBF, we used the anchor-away technique*(37)*. A rapamycin-resistant strain background that contained a mutated *TOR1 (tor1-1)* and deleted *FPR1 (fpr1*Δ) was used and the anchor *(SWI4* or *SWI6)* was C-terminally tagged with *FKBP12* (human 12 kDa FK506 binding protein) and an extra copy of C-terminally tagged *KIN28* was C-terminally tagged with *FRB* (11 kDa, FKBP12-rapamycin-binding domain of human mTOR) and expressed ectopically as a fusion with its cyclin partner *CCL1* from the *CLN3* promoter *(CLN3pr-CCL1-L-KIN28-FRB)*. To induce protein binding, cells were cultured at 30°C in SCD media with1 μg/ml rapamycin, or DMSO as a control.

### Cell size measurements

Cell volume was measured using a Beckman Coulter Z2 counter (Beckman Coulter). Log-phase cultures at OD_600_ 0.2-0.3 were briefly sonicated, and then 100-150 µL was diluted into 10 mL of Isoton II diluent (Beckman Coulter #8546719), and 40,000-50,000 cells were measured per sample. Particles below 10 fL and over 300 fL in volume were excluded from analysis.

### Immunoblotting

Protein lysates were taken in urea lysis buffer as previously described*(31)*. Protein lysates were separated on tris-glycine or tris-acetate SDS-PAGE gels and transferred to a nitrocellulose membrane using the iBlot 2 dry blotting system (Invitrogen IB21001). The following primary antibodies where used for western-blotting at 1/1000 dilution: anti-Rpb1-CTD clone 8wG16 (mouse, monoclonal, Abcam), anti-Rpb3 clone 1Y26 (mouse, monoclonal, BioLegend), anti-Rpb1-S2-P clone 3E10 (rat, monoclonal, Millipore), anti-Rpb1-S5-P clone H14 (mouse, monoclonal, BioLegend), anti-Rpb1-S5-P clone 3E8 (rat, monoclonal, Millipore), anti-Rpb1-S7-P clone 3E12 (rat, monoclonal, Millipore), anti-Cdc28 clone yC-20 (goat, polyclonal, Santa Cruz), P-cdc2 (T161) Rabbit antibody (Cell Signaling Technology) or anti-FLAG; clone M2 (mouse, monoclonal, SIGMA). Primary antibodies were detected using the following fluorescently labeled secondary antibodies at 1/10,000 dilution: IRDye 680LT Goat anti-Mouse, IRDye 680LT Goat anti-Rat, IRDye 800CW Goat anti-Rat, IRDye 800CW Goat anti-Mouse (Licor), Alexa Fluor 680 Donkey anti-Mouse, Alexa Fluor 680 donkey anti-Rabbit and Alexa Fluor 790 Goat anti-Rabbit (Invitrogen by Thermo Fisher Scientific). Membranes were then imaged on a LI-COR Odyssey CLx.

### Immunoblot quantifications

Band intensities where quantified using the LI-COR ImageStudio Lite software. For the quantifications in Fig. 3L, the Rpb1-CTD-S5-P signal was normalized for loading to Rpb3. To calculate the Kin28-dependent fraction of Rpb1-CTD-S5-P, the difference between before (t=0) and after CMK addition (t=20) in the WT *(i*.*e*., not *cln3*Δ) background was calculated and divided through its value at t=0. Then, to calculate the Cln3-dependent Rpb1-CTD-S5-P fraction, the difference between the *cln3*Δ and *CLN3* wild type background after CMK addition (t=20) was calculated and divided through its value at t=0 in the *CLN3* wild type background. The mean and S.E.M. were calculated from two independent biological replicates.

### Spot viability assay

Plate spot assays show cell growth on YP plate with 2% glucose or 2% galactose from a series of culture dilutions (2x, 10x, 5x, and 10x) from an initial amount of 106 cells. Plates were incubated at 30°C and photographed at least 40h later.

### Microscopy

Cells were grown in a CellASIC microfluidic device as previously described*(5)*. For the experiment in Fig. 2E cells were grown in SC medium with 2% galactose for 2h before media was replaced with SC 2% glucose to turn off the expression of endogenous *GAL1pr-CDK1*. Phase-contrast images were acquired every 3 minutes after media shift, and multiple fields of view were followed simultaneously. For full movies see supporting material Movies S1-4.

### ChIP-seq experiments

Cells expressing Cln3-V5, Swi4-V5, Swi6-V5 or GFP-NLS-V5 were grown in SC media with 2% glycerol 1% ethanol. 250ml of cells at OD ∼0.6 were fixed with 1% formaldehyde (30 minutes) and quenched with 0.125 M glycine (5 minutes). Fixed cells were washed twice in cold PBS, pelleted, snap-frozen and stored at −80°C. Cell lysis and ChIP reactions were performed as previously described*(52)* with minor modifications. Pellets were lysed in 300 µL FA lysis buffer (50 mM HEPES–KOH pH 8.0, 150 mM NaCl, 1 mM EDTA, 1% Triton X-100, 0.1% sodium deoxycholate, 1 mM PMSF, Roche protease inhibitor) with ∼1 mL ceramic beads on a Fastprep-24 (MP Biomedicals). The entire lysate was then collected and adjusted to 1 mL before sonication with a 1/8’ microtip on a Q500 sonicator (Qsonica) for 16 minutes (10 seconds on, 20 seconds off). The sample tube was held suspended in a −20°C 80% ethanol bath to prevent sample heating during sonication. Cell debris was then pelleted and the supernatant retained for ChIP. For each ChIP reaction, 30 µL Protein G Dynabeads (Invitrogen) were blocked (PBS + 0.5% BSA), prebound with 5-10 µL anti-V5 antibody (SV5-Pk1, BioRad Cat# MCA1360G) and washed once with PBS before incubation with supernatant (4°C, overnight). Dynabeads were then washed (5 minutes per wash) twice in FA lysis buffer, twice in high-salt FA lysis buffer (50 mM HepesKOH pH 8.0, 500 mM NaCl, 1 mM EDTA, 1% Triton X-100, 0.1% sodium deoxycholate, 1 mM PMSF), twice in ChIP wash buffer (10 mM TrisHCl pH 7.5, 0.25 M LiCl, 0.5% NP-40, 0.5% sodium deoxycholate, 1 mM EDTA, 1 mM PMSF) and once in TE wash buffer (10 mM TrisHCl pH 7.5, 1 mM EDTA, 50 mM NaCl). DNA was eluted in ChIP elution buffer (50 mM TrisHCl pH 7.5, 10 mM EDTA, 1% SDS) at 65°C for 20 minutes. Eluted DNA was incubated to reverse crosslinks (65°C, 5hr), before treatment with RNAse A (37°C, 1 hour) and then Proteinase K (65°C, 2 hours). DNA was purified using the ChIP DNA Clean & Concentrator kit (Zymo Research). Indexed sequencing libraries were generated using the NEBNext Ultra II DNA Library Prep kit (NEB Cat # E7645), pooled and sequenced on an Illumina HiSeq instrument as paired end reads (Novogene, CA).

### ChIP-seq analysis

Data processing was performed in Galaxy (https://usegalaxy.org/). Reads were trimmed to 36bp using *Cutadapt* and then aligned to the *S. cerevisiae* genome (SacCer3) using *Bowtie2*. RPKM normalized Bigwig files were generated using *bamCoverage* (bin size =10bp, paired end reads extended, smoothing = 100bp) and used for track display with *Integrative Genome Viewer*. BAM files were filtered to remove duplicate and low-quality reads with *BAM filter* before peak calling with *MACS2* using GFP-NLS-V5 as the control (genome size = 12000000, bandwidth = 200). Cln3 peaks were defined as regions in which peaks were identified in both Cln3-V5 ChIP replicates (n=58) and SBF peaks were defined as regions in which peaks where identified in Swi4-V5 and Swi6-V5 ChIP experiments (n=311). SGD genes where downloaded from *UCSC Main* in BED format and promoters were defined as 1kb upstream of the ORF start. Table S3 contains the list of 85 promoters which overlapped with the 58 Cln3 peaks.

### Protein expression and purification

Full-length N-terminally glutathione S-transferase-tagged (GST-tagged) proteins were expressed in the *E. coli* strain BL21 and purified by glutathione-agarose affinity chromatography (Sigma-Aldrich Cat #G4510) and eluted using elution buffer (50mM Tris pH 8.0, 100mM KOAc, 25mM MgOAc, 10% glycerol, 15mM glutathione). N-terminally 6His-tagged recombinant substrates were expressed in the *E. coli* strain BL21 and the purification was performed using cobalt affinity chromatography. Proteins were eluted using buffer containing 50 mM HEPES pH 7.4, 150mM NaCl, 10% glycerol, and 200mM imidazole. Histone H1 protein, which was used as a general substrate for Cdk1, was purchased from EMD Millipore (Cat #14-155). GST-4CTD fusion proteins containing a GST-tag and 4 repeats of the Rpb1 unstructured CTD consensus repeat or repeats with substitutions in specific residues were expressed and purified as described above.

All cyclin-Cdk1 fusion complexes were purified from budding yeast cells using a 3X FLAG affinity purification method modified from a previous protocol used for HA-tag purification*(53)*. Briefly, N-terminally tagged cyclin-Cdk1 fusions were cloned into a pRS425 vector using a glycine-serine linker*(24)* and overexpressed from the *GAL1* budding yeast promoter. The use of a glycine-serine rich linker was a key step in Cln3-L-Cdk1 purification as the Cln3 protein had notably lower affinity towards Cdk1 in high salt conditions than other *S. cerevisiae* cyclins. The overexpressed 3X FLAG-tagged cyclin-Cdk1 complexes were then purified by immunoaffinity chromatography using ANTI-FLAG M2 affinity agarose beads (Sigma-Aldrich Cat #A2220) and eluted with 0.2 mg/mL 3X FLAG peptide (Sigma-Aldrich Cat #F4799). We note that similar cyclin-Cdk fusions have previously been able to restore wild-type function *in vivo(23)*. Cks1 was purified as described previously*(54)* and added separately to Cdk1 enzyme complexes in all phosphorylation assays.

### *In vitro* phosphorylation assays

For all phosphorylation assays, equal amounts of substrate and purified kinase complexes were used. Substrate concentrations were kept in the range of 1-5 μM for different experiments but did not vary within any experiment. Reaction aliquots were taken at two time points (if not stated otherwise, at the 8- and 16-minute time points) and the reaction was stopped with SDS-PAGE sample buffer. The basal composition of the assay mixture contained 50 mM HEPES pH 7.4, 150 mM NaCl, 5 mM MgCl2, 0.2 mg/ml 3X FLAG peptide, 6% glycerol, 3 mM EGTA, 0.2 mg/ml BSA, and 500 µM ATP (with 2 μCi of [γ-32P] ATP added per reaction; PerkinElmer BLU502Z250UC). Phosphorylated proteins were separated on 10% SDS-PAGE gels unless stated otherwise. In the case of GST-4CTD model substrates, 12% SDS-PAGE gels were used, while in the case of full length Rpb1 and Rpb1ΔC, 6% SDS-PAGE gels were used. Phosphorylation of substrate proteins was visualized using autoradiography (Typhoon 9210; GE Healthcare Life Sciences). Autoradiographs were quantified with the ImageQuant TL Software (GE Healthcare Life Sciences).

### Baculovirus-based Cyclin-Cdk1-Cks1 complex expression and purification

Baculovirus-based expression of yeast Cyclin-Cdk1-Cks1 complexes was performed following *(4)*, as described in *(55)*. Briefly, Sf9 insect cells (gift of Tim Stearns) were grown to confluence in a 75 cm^2^ culture flask in Sf-900 II SFM media (Thermo-Fisher, Waltham, MA) and infected with 3 mL, 1.5 mL, and 1.5 mL, respectively, of GST-Cdk1, Cln3, and Cks1 baculovirus stocks (Gift of Mike Tyers). For the no-Cln3 control, Cln3 virus was omitted. After 40 hours of infection, cells were harvested, pelleted, and frozen before being used for purification as follows. Frozen insect cell pellets were thawed and resuspended in 2 mL of ice-cold lysis buffer (50 mM HEPES pH 7.2, 150 mM NaCl, 5 mM EDTA, 0.1% v/v NP-40, 10% v/v glycerol) supplemented with protease inhibitors (1 µg/mL Pepstatin A, 1 µg/mL Leupeptin, >2 KIU/mL Aprotinin, 1 mM Benzamidine, 1 µg/mL Bestatin, 2 mM PMSF) and phosphatase inhibitors (50 mM NaF, 1 mM Sodium orthovanadate, 80 mM Beta-glycerophosphate). Resuspended cells were incubated for 30 minutes on ice before being centrifuged for 10 minutes at 18,000 x g at 4°C. The supernatant was transferred to a new tube and centrifuged for 20 minutes at 28,000 x g at 4°C. The cleared lysate was mixed with 200 µL of glutathione agarose beads (Sigma-Aldrich, St. Louis, MO) and turned end-over-end for 1.5 hours at 4°C before being loaded into a gravity column. The column was washed with 60 volumes of lysis. Before elution, the beads were resuspended in 600 µL lysis buffer, and 100 µL of the resulting 25% slurry was removed for use in on-bead kinase reactions. The beads were allowed to re-settle, and protein was eluted from the resulting column twice, using 175 µL of lysis buffer + 25 mM glutathione each time. The beads were incubated in elution buffer for 1 hour before the first elution and 1.5 hours before the second elution. Eluates were aliquoted and flash frozen.

### Kinase assays using glutathione-agarose-bound kinase activity

For each reaction using bead-bound kinase activity, 20 µL of the 25% glutathione-agarose slurry sampled above was centrifuged to give a 5 µL bead pellet. The supernatant was removed, and 5 µL of 4x kinase buffer supplemented with radiolabeled ATP, 0.8 mg/mL BSA, and 100 nM Cks1 was added. The resulting 10 µL of 50% slurry was then mixed with 10 µL of substrate solution, as in the kinase assays described above. Tubes were agitated at regular intervals during kinase reactions to keep the beads in suspension, and samples were removed using cut pipet tips, to ensure bead capture.

## Acknowledgements

We thank Tim Stearns for Sf9 cells, Mike Tyers for baculovirus stocks, Jennifer Ewald, Kurt Schmoller, David Morgan, Steve Hahn, and Steve Buratowski for yeast strains. We thank Mart Loog, Jim Ferrell, Fred Cross, Peter Pryciak, Aseem Ansari, David Morgan, and Skotheim lab members for constructive feedback. We thank Ben Reyes Topacio for help with enzyme purifications. This work was supported by the NIH (GM092925 and GM115479), the HHMI-Simons (JMS Faculty Scholars Program), the HFSP (postdoctoral fellowship to MK), and the Life Sciences Research Foundation (Simons Foundation Fellowship to MS).

---

**Fig. S1.**
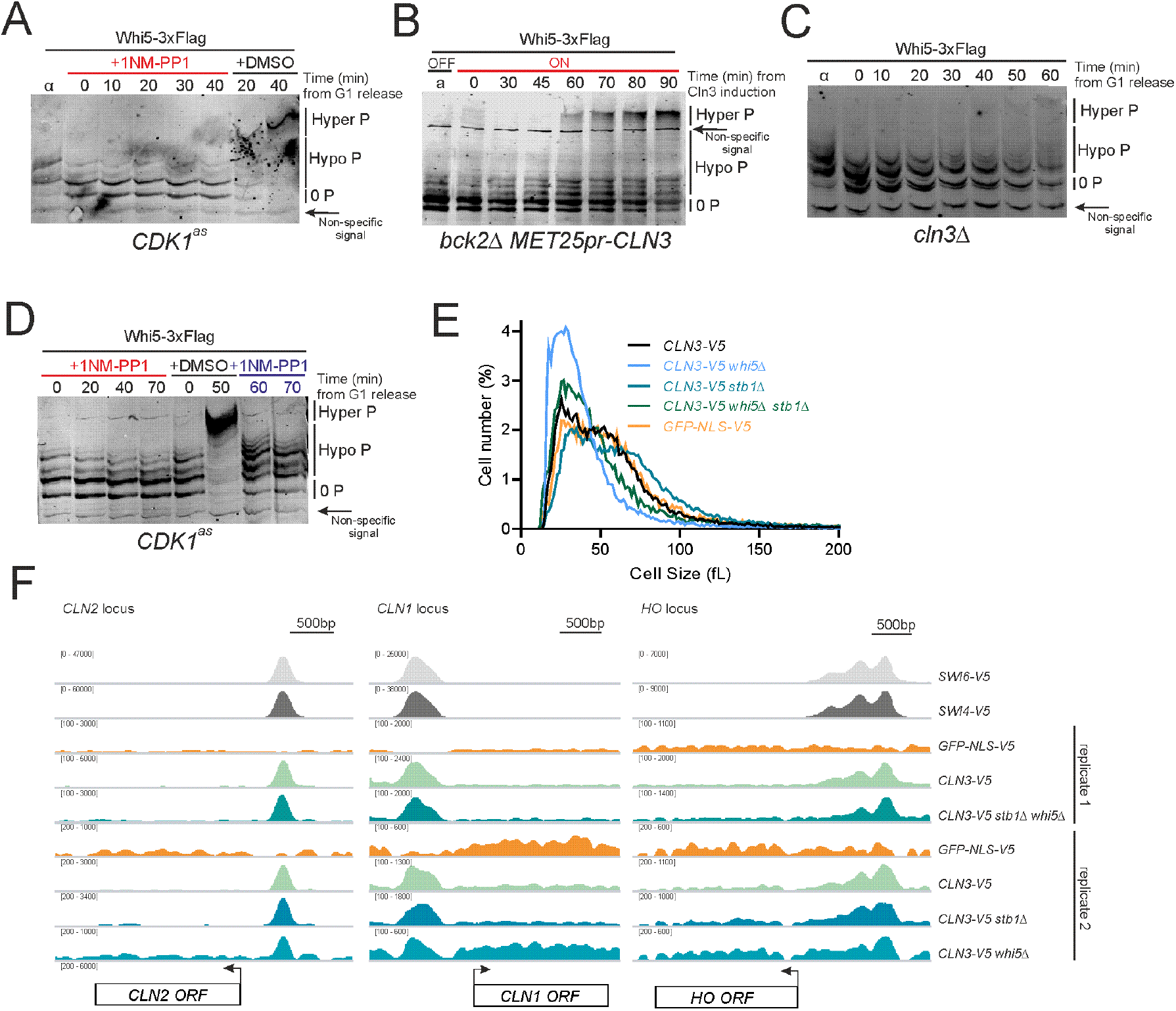
(**A**) to (**D**) Phos-tag immunoblots measuring distinct hypo- and hyper-phosphorylated isoforms of Whi5 C-terminally tagged with 3 Flag epitopes expressed from the *WHI5* promoter (Whi5-3xFlag). (**A**) Whi5-3xFlag phosphorylation time series in the ATP analog sensitive *CDK1*^*as*^ strain. Cells were released from pheromone-induced G1 arrest into media with DMSO or 10µM of the ATP analog 1NM-PP1. (**B**) Whi5-3xFlag phosphorylation time series in a *bck2*Δ *MET25pr-CLN3* strain. Cells arrested in G1 in the presence of methionine and then were released into media without methionine. (**C**) Whi5-3xFlag phosphorylation time series after *cln3*Δ cells were released from a pheromone-induced G1 arrest. (**D**) Whi5-3xFlag phosphorylation time series in a *CDK1*^*as*^ strain. Cells were released from a pheromone-induced G1 arrest and then DMSO or 10µM 1NM-PP1 were added at 0 minutes when Whi5 was hypophosphorylated. In the case where DMSO was initially added, we then also added 10µM 1NM-PP1 at 50 minutes when Whi5 is hyperphosphorylated. (**E**) Cell size distributions measured by Coulter counter for the strains used in the Cln3-V5 ChIP experiments used to generate the data for Fig. 1E and fig. S1F. All strains were grown on synthetic complete media +2% glycerol +1% ethanol. (**F**) ChIP-seq signal at three example SBF regulated genes: *CLN2, CLN1* and *HO. CLN3, SWI4* or *SWI6* were tagged at the endogenous loci with the V5 epitope and anti-V5 ChIP-seq was performed in the indicated genotypes as described in the methods. A subset of these data are also presented in Fig. 1E.

**Fig. S2.**
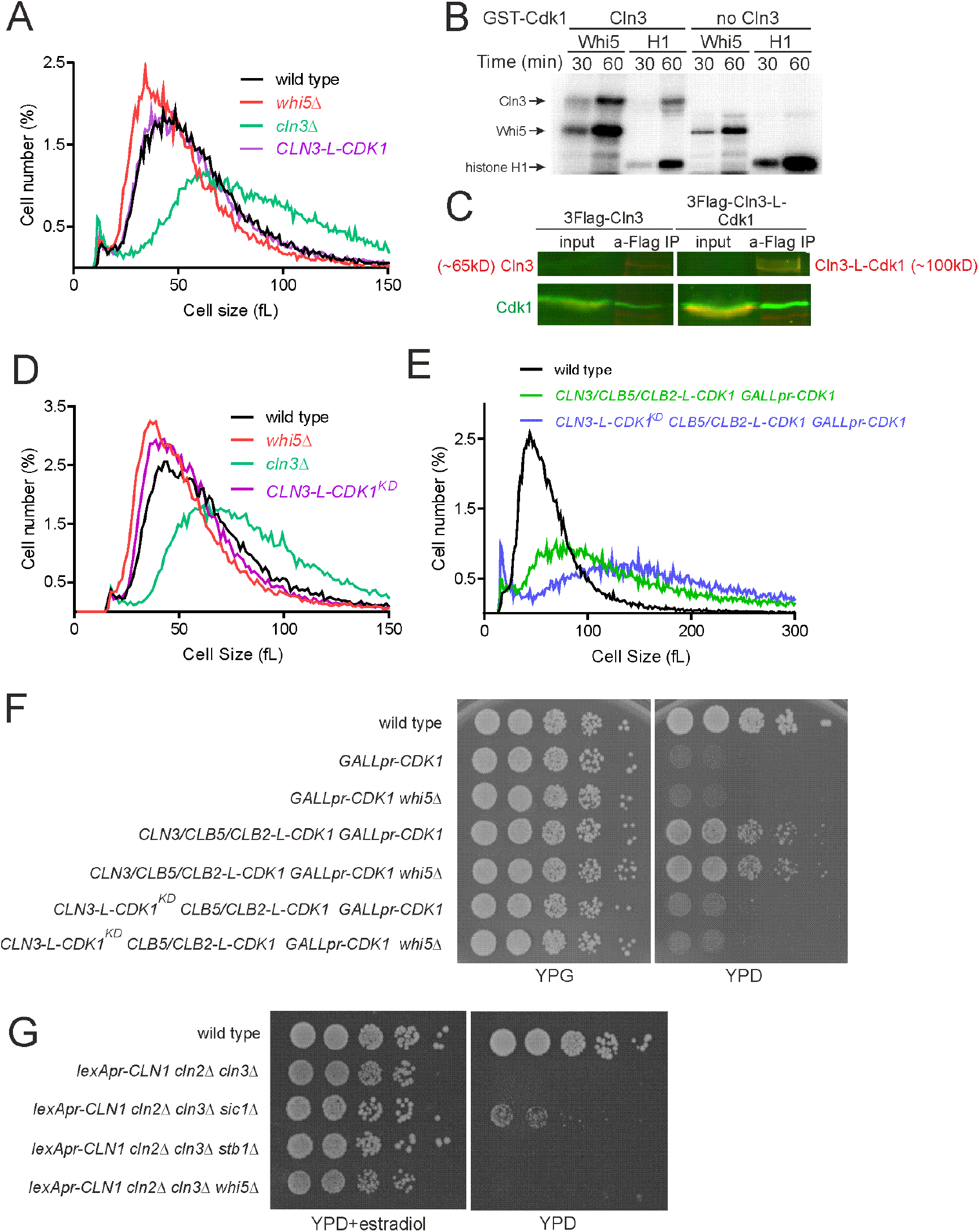
(**A**) Cell size distributions measured by Coulter counter for the indicated genotypes. Cells were grown on synthetic complete media +2% glucose. (**B**) Autoradiographs of *in vitro* kinase assays using substrates Whi5 and histone H1. Kinase activity was purified from Sf9 insect cells expressing Cln3, Cks1 and Cdk1-GST (left) or only Cks1 and Cdk1-GST (right). The activity previously reported in *(4)* to be due to Cln3-Cdk1 in was present in the control purification without Cln3 expression as seen on the right-hand side panel (see methods). (**C**) Immunoblots of samples following immunoprecipitation with 3xFlag-Cln3 or 3xFlag-Cln3-L-Cdk1^KD^, both expressed from the *GAL1* promoter. Both 3xFlag-Cln3 and 3xFlag-Cln3-L-Cdk1^KD^ co-immunoprecipitated with endogenous Cdk1. (**D**) Cell size distributions, measured by Coulter counter, for the indicated genotypes. Cells were grown on synthetic complete media +2% glucose. (**E**) Cell size distributions, measured by Coulter counter, for the indicated genotypes. Cells were grown on synthetic complete media +2% galactose before adding 2% glucose to repress *GALL* promoter-dependent expression of Cdk1. Cell size was measured 12 hours after *GALLpr* repression. (**F**) Spot viability assays of WT and strains with *GALLpr*-dependent expression of Cdk1 on YPG *(GALLpr* ON) or YPD *(GALLpr* OFF). Adding three cyclin-Cdk1 fusion genes rescues *GALLpr-CDK1* repression. However, if *CLN3-CDK1* is replaced with a kinase dead *CLN3-CDK1*^*KD*^ fusion, cells are not viable even if *WHI5* is deleted. (**G**) Spot viability assays of WT and strains where G1 is driven exclusively by the hormone responsive promoter *(LexApr)* expressing *CLN1* on YPD + 50nM Beta-estradiol *(LexApr* ON) or YPD *(LexApr* OFF). Deletion of *SIC1*, but not *whi5*Δ or *stb1*Δ restores viability.

**Fig. S3.**
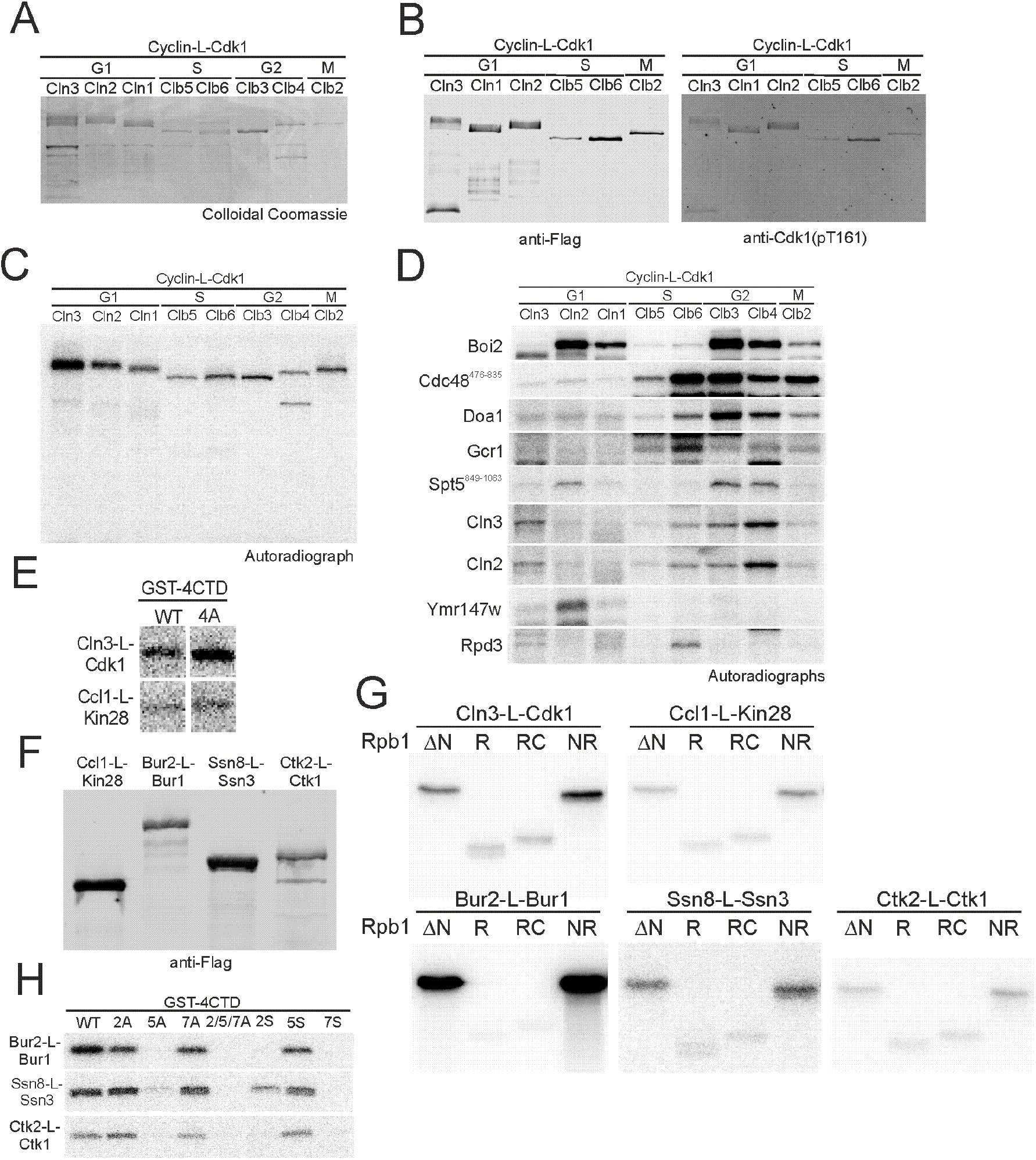
(**A**) Colloidal Coomassie-stained SDS gel of all purified cyclin-L-Cdk1 preparations used in this study. (**B**) Immunoblots to determine the levels (anti-FLAG) and activating phosphorylation (anti-Cdk1 pT169) of the indicated purified cyclin-L-Cdk1 complexes used in this study. (**C**) Autoradiographs of *in vitro* kinase assays using the indicated purified cyclin-L-Cdk1 complexes without any substrate protein. The signal is due to autophosphorylation. (**D**) Autoradiographs of *in vitro* kinase assays using equal amounts of the denoted cyclin-L-Cdk1 and candidate Cln3-Cdk1 substrates. (**E**) Autoradiographs of *in vitro* kinase assays using Cln3-L-Cdk1 and Ccl1-L-Kin28 to phosphorylate a synthetic substrate containing 4 CTD repeats or 4 CTD repeats with threonine 4 mutated to alanine. (**F**) Immunoblots to determine the levels (anti-FLAG) of purified transcriptional cyclin-L-Cdk complexes used in this study. (**G**) Autoradiographs of *in vitro* kinase assays with transcriptional cyclin-L-Cdk complexes phosphorylating Rpb1ΔN and a series of Rpb1ΔN truncations. See Fig. 3E for details of truncations. (**H**) Autoradiographs of *in vitro* kinase assays using Cln3-L-Cdk1 or transcriptional cyclin-L-Cdks to phosphorylate WT or mutant versions of 4 CTD repeats. See Fig. 3F for details of CTD repeat variants.

**Fig. S4.**
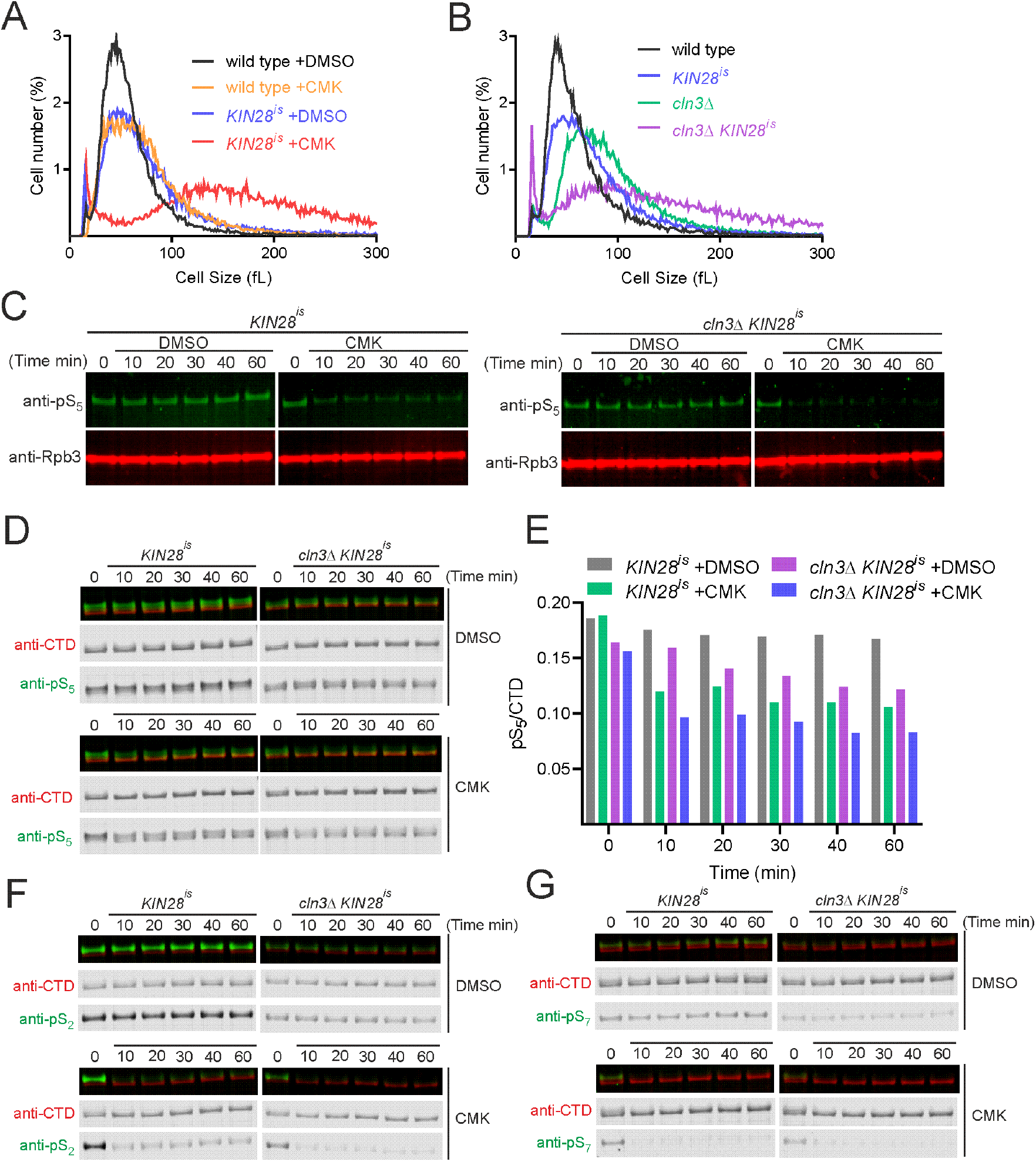
(**A**) Cell size distributions measured by Coulter counter for the indicated genotypes. All strains were grown on synthetic complete media +2% glucose with 5µM CMK or DMSO. (**B**) Cell size distributions measured by Coulter counter for the indicated genotypes. All strains were grown on synthetic complete media +2% glucose. (**C**) to (**G**) Immunoblots and quantifications of different phosphorylated forms of Rpb1 in *KIN28*^*is*^ or *KIN28*^*is*^ *cln3*Δ strains at the indicated timepoints after release from G1 pheromone arrest into DMSO or 5µM CMK. *Kin28*^*is*^ contains an active site mutant rendering it sensitive to covalent inhibition by the small molecule CMK. (**C**) Immunoblots of total cellular phosphorylated Rpb1-CTD S_5_ (H14 antibody) and total cellular RNA polymerase II (Rpb3) after release from G1 pheromone arrest into DMSO or 5µM CMK. A subset of these data are also presented in Fig. 3K. (**D**) Immunoblots of total cellular phosphorylated Rpb1-CTD S_5_ (3E8 antibody) and Rpb1-CTD (8wG16 antibody) after release from G1 pheromone arrest into DMSO or 5µM CMK. (**E**) Quantification of immunoblots in (D): total cellular phosphorylated CTD S_5_ (3E8 antibody) normalized to Rpb1-CTD (8wG16 antibody). (**F**) Immunoblots of total cellular phosphorylated Rpb1-CTD S_2_ (3E10 antibody) and Rpb1-CTD (8wG16 antibody) after release from G1 pheromone arrest into DMSO or 5µM CMK. (**G**) Immunoblots of total cellular phosphorylated Rpb1-CTD S_7_ (3E12 antibody) and Rpb1-CTD (8wG16 antibody) after release from G1 pheromone arrest into DMSO or 5µM CMK.

**Fig. S5.**
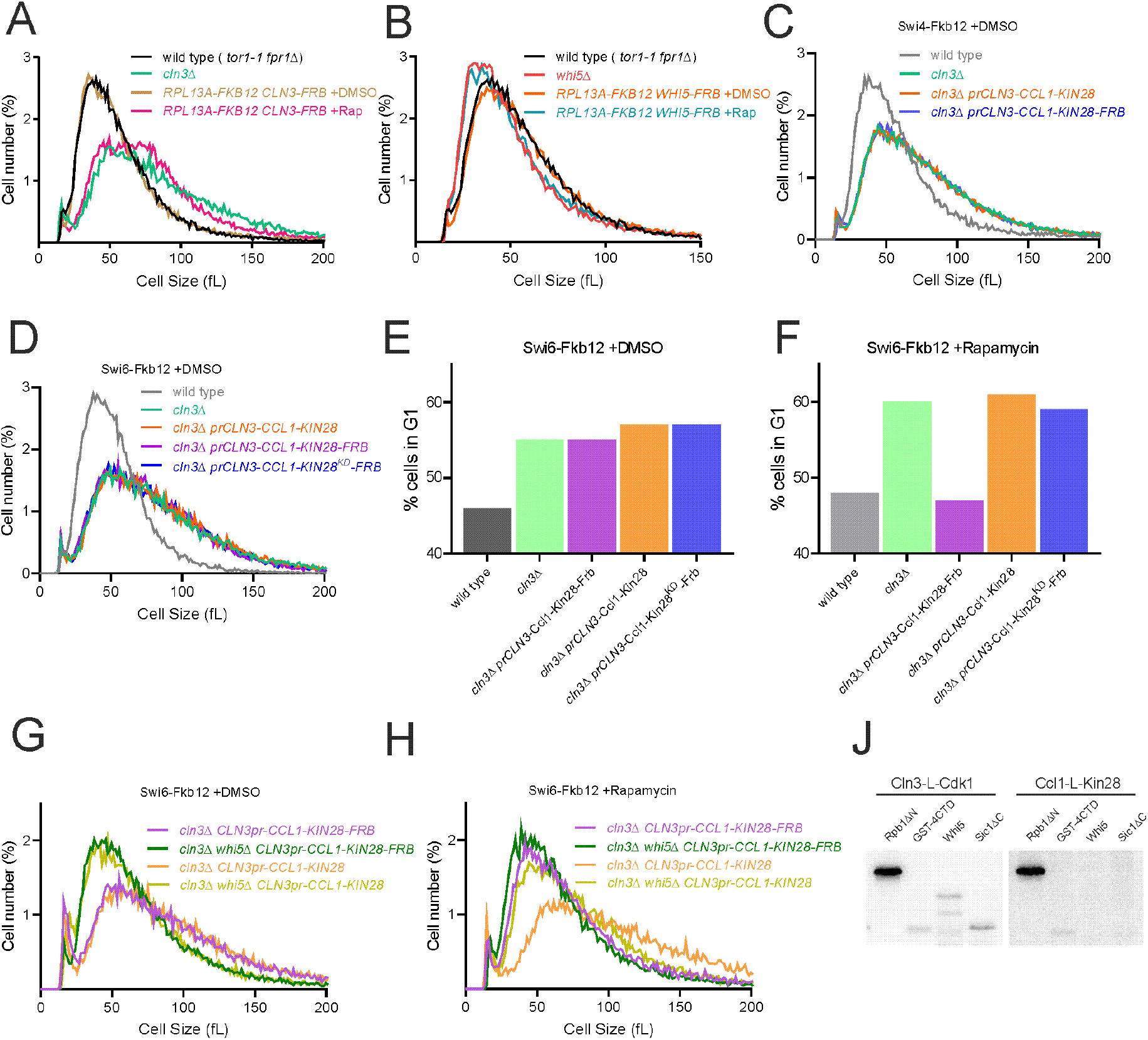
(**A**) to (**B**) Cell size distributions measured by Coulter counter for the indicated genotypes. All strains were grown on synthetic complete media +2% glucose. 1µg/ml rapamycin or DMSO was added ∼200 minutes before cell size measurements. (**C**) to (**H**) Conditional recruitment of Ccl1-L-Kin28 to SBF using the rapamycin inducible binding system. Ccl1-L-Kin28 fusion proteins were expressed from a genomically integrated copy of the *CLN3* promoter. Ccl1-L-Kin28-FRB was recruited to SBF via its Swi4 (C&G) or Swi6 (D to F & H) subunits upon rapamycin treatment. Ccl1-L-Kin28 lacking FRB is not recruited. All strains were grown on synthetic complete media +2% glucose with 1µg/ml rapamycin or DMSO. (**C**) to (**D**) Cell size distributions measured by Coulter counter for the indicated genotypes. (**E**) to (**F**) Quantification of flow cytometry analysis of DNA content of the cells in (D) and Fig. 5D. Samples were collected from the same cultures at the same time for both cell size and DNA content measurements. (**G**) to (**H**) Cell size distributions measured using a Coulter counter for the indicated genotypes. (**J**) Autoradiographs of *in vitro* phosphorylation of Rpb1ΔN, GST-4CTD, Whi5 or Sic1ΔC by Cln3-L-Cdk1 or Ccl1-L-Kin28. Ccl1-L-Kin28 is not able to phosphorylate Whi5.

